# Glial *betaPix* is essential for blood vessel development in the zebrafish brain

**DOI:** 10.1101/2025.04.08.647738

**Authors:** ShihChing Chiu, Qinchao Zhou, Chenglu Xiao, Linlu Bai, Xiaojun Zhu, Wanqiu Ding, Jing-Wei Xiong

**Author notes:** Corresponding authors: WD, XZ, and JWX. Lead Contact: Jing-Wei Xiong, Ph.D.

## Abstract

The formation of blood-brain barrier and vascular integrity depends on the coordinative development of different cell types in the brain. Previous studies have shown that zebrafish *bubblehead* (*bbh)* mutant, which has mutation in the *betaPix* locus, develops spontaneous intracerebral hemorrhage during early development. However, it remains unclear in which brain cells *betaPix* may function. Here, we established a highly efficient conditional knockout method in zebrafish by using homology-directed repair (HDR)-mediated knockin and knockout technology, and generated *betaPix* conditional trap (*betaPix*^*ct*^) allele in zebrafish. We found that *betaPix* in glia, but neither neurons, endothelial cells, nor pericytes, was critical for glial and vascular development and integrity, thus contributing to the formation of blood-brain barrier. Single-cell transcriptome profiling revealed that microtubule aggregation signaling stathmins and pro-angiogenic transcription factors *Zfhx3/4* were down-regulated in glial and neuronal progenitors, and further genetic analysis suggested that betaPix may act upstream on the PAK1-Stathmin and Zfhx3/4-Vegfaa signaling to regulate glia migration and angiogenesis. Therefore, this work reveals that glial *betaPix* plays an important role in brain vascular development in zebrafish embryos and possibly human cells.

## Introduction

Intracerebral hemorrhage (ICH) is a life-threatening stroke type with the worst prognosis and few proven clinical treatments. Most patients who survive intracerebral hemorrhage end up with disabilities and are at risk for recurrency, cognitive decline, and systemic vascular issues, making this condition particularly significant among neurological disorders^1,2^. Several rodent models have been used for modeling ICH, including autologous blood injection, collagenase injection, thrombin injection, and micro-balloon inflation techniques^3^. Zebrafish (Danio rerio) exhibit a closed circulatory system and regulatory pathways that are highly conserved among vertebrates ^4^ with relatively cheap and easy to maintain disease models. Over the past decades, varieties of zebrafish mutants that spontaneously developed brain hemorrhage have been generated in mutagenesis screens, such as *redhead* ^5^, *reddish* ^6,7^ and *bubblehead (bbh)*^8,9^. Elucidating mutated gene function in these zebrafish models enables us to gain insights into the genetic basis and pathophysiological mechanisms of ICH development. The *bbh* mutant was identified independently in two large-scale ethylnitrosourea-induced mutagenesis screens ^8,9^ and positional cloning revealed that p21-activated kinase (Pak)-interacting exchange factor beta *(betaPix)* gene was mutated in *bbh* mutants^10^. *betaPix* contains SH3 domain that binds group I PAKs, and Dbl homology (DH) and pleckstrin homology (PH) domains that function as a RhoGEF to interact with Rac/Cdc42 small GTPases ^11^, thus participating in multiple cellular pathways to regulate cell polarity, adhesion and migration ^12^.

Various studies have reported that brain vessels are highly supported by neurons and glial cells during development and regeneration ^13,14^ Glial-specific *betaPix* have been proposed to interact with *αv/β8* integrin and the Band 4.1s in glia, which link to multiple intracellular signaling effectors such as *Wnt7a* and *Wnt7b* ^15^, but this hypothesis was not experimentally addressed ^16^. To this end, *betaPix* has been shown to interact with *αvβ8* integrins and mediate focal adhesion formation and thus modulate cerebral vascular stability in *bbh* zebrafish mutant ^17^. Integrins are the main molecular link between cells and the extracellular matrix (ECM) which serves as scaffolds in the perivascular space. Based on a chemical suppressor screening of *bbh* hemorrhages, we have previously reported that a small molecule called miconazole downregulated the pErk-matrix metalloproteinase 9 (Mmp9) signaling to reduce ECM degradation, thus improving vascular integrity ^18^. However, how *betaPix* cell-type specifically contributes to vascular integrity and maturation remains unclear. Here, we generated *betaPix* conditional trap *(betaPix*^*ct*^) alleles by using a CRISPR/HDR-based Zwitch method, and found that *betaPix* acted mainly in glia to regulate glial and vascular development and integrity *via* Stathmin and Zfhx3/4 signaling. Therefore, this work provides the first experimental evidence that *betaPix* acts in glia for vascular integrity development, and its functional conservation between zebrafish and human cells may further guide us to decipher the genetic basis of ICH in the future.

## Results

### Generating *betaPix* conditional trap (*betaPix^ct^*) allele by an HDR-mediated knockin and knockout method

To study *betaPix* function in cell-specific manner, we utilized a donor vector with an invertible gene trap cassette *via* HDR (Zwitch) ^19–21^ for generating *betaPix* conditional trap *(betaPix*^*ct*^) alleles. Zwitch consists of a splice acceptor conjugated to a red fluorescent protein *via* a 2A self-cleaving peptide, flanked by two LoxP sites and two Lox 5171 sites in reciprocal orientations. Outside this conditional trap cassette, we included a lens-specific enhanced green fluorescence protein *α*-crystallin:EGFP for screening knockout founders, as well as inserted both left and right arms for targeting homologous sequences on the genome. We modified the Zwitch vector by adding a glycine-serine-glycine (GSG) spacer in front of the 2A peptide to enhance cleavage efficiency, and adding universal guide RNA (UgRNA)^22^ target sequences outside the left and right homologous arms for linearizing donor vector (Figures 1A and S1A). We chose CRISPR/Cas9 system to generate double-strand breaks (DSBs) on intron 5 of the *betaPix* locus, and chose the most efficient guide RNAs based on their efficiency to induce CRISPR indels. We used either long arms such as ~1000 bp or short arms such as 24 bp that located upstream and downstream the guide RNA sites. We then co-injected the donor vectors with targeting guide RNA, universal guide RNA, and Cas9 mRNA into one-cell-stage embryos. At 4 days post-fertilization (dpf), we selected *α*-crystallin-EGFP-positive embryos (F_0_) for raising to adulthood, and pre-selected F_0_ founders by examining inheritable EGFP reporter expression of F_1_ embryos. To confirm the correct homologous recombination in the *betaPix* locus, we identified potential founders by genomic PCR to examine Zwitch insertion in the *betaPix* locus. Precise knock-in genotypes were further verified using Sanger sequencing (Figure 1B, 1C and S1B). Among 184 adult F_0_ founders with *α*-crystallin-EGFP expression, we found 84 founders had EGFP-positive F_1_ embryos, and 7 founders had expected PCR fragments around both 5’ and 3’arms, which two founders had correct insertions in the *betaPix* locus by Sanger sequencing (Figure S1C), thus achieving ~1% efficiency on generating conditional alleles. Briefly, we established an efficient method for generating conditional knockin/knockout mutants in general, and generated a conditional gene trap line Ki*(betaPix*:Zwitch) in the *betaPix* locus in particular, which is referred to as *betaPix*^*ct*^ hereafter.

**Figure 1.**
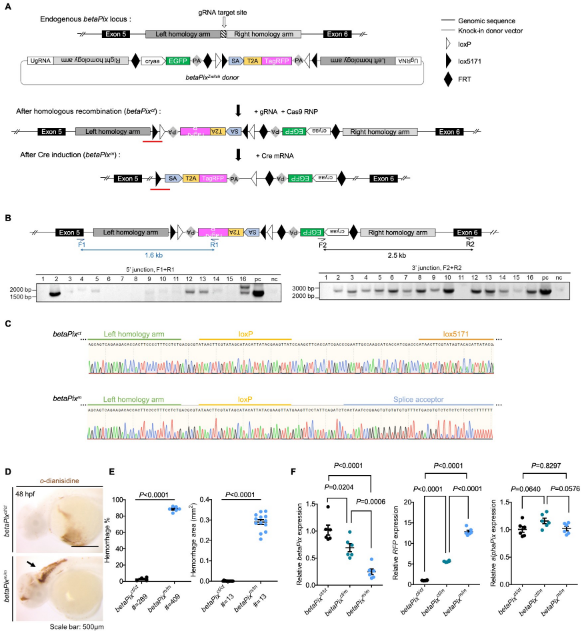
Generation of *betaPix* conditional trap *(betaPix*^*ct*^) allele by an homologous recombination (HDR)-mediated knock-in method. (A) Schematic diagram illustrates the HDR-mediated *Zwitch* strategy for generating zebrafish knock-in allele at the *betaPix* locus. (B) Genomic PCR analysis of the F_1_ embryos confirming the right *Zwitch* insertions. (C) Sanger sequencing confirming the junction of *betaPix*^*ct*^ (after HDR-mediated insertion) or *betaPix*^*m*^ (after Cre-mediated inversion) that are highlighted by RED lines in (A). (D) Representative stereomicroscopy images of erythrocytes stained with *o*-dianisidine in *betaPix*^*ct/ct*^ and *betaPix*^*m/m*^ embryos at 48 hpf. Brain hemorrhages, indicated with arrow, in *Cre* mRNA-injected embryos *(betaPix*^*m/m*^). Lateral views, anterior to the left. (E) Quantification of hemorrhagic parameters in (D). Left panel showing hemorrhage percentages, with independent experiments as dots. Right panel showing hemorrhage areas, with each dot representing one embryo, # represents the numbers of embryos scored for each analysis, and three or more individual experiments conducted. Data are presented in mean ± SEM; unpaired Student’s t test with individual *P* values mentioned in the figure. (F) qRT-PCR analysis showing the expression of *betaPix, RFP*, and *alphaPix* in *betaPix*^*ct/ct*^, *betaPix*^*ct/m*^, and *betaPix*^*m/m*^ embryos at 48 hpf. Each dot represents one embryo. Data are presented in mean ± SEM; one-way ANOVA analysis with Dunnett’s test, individual *P* values mentioned in the figure. cryaa, α A-crystallin; PA, polyadenylation signal; SA, splice acceptor; T2A, T2A self-cleaving peptide. Individual scale bars indicated in the figure.

A previous work has shown that *bubblehead* (*bbh*^*fn40a*^) mutant has a global reduction in *betaPix* transcripts, and *bbh*^*m292*^ mutant has a hypomorphic mutation in *betaPix*, thus establishing that *betaPix* is responsible for *bubblehead* mutant phenotypes^10^. *bbh*^*fn40a*^ mutants developed cerebral hemorrhages phenotype between 36 to 52 hpf (Figure S1D and S1E). To characterize the *betaPix*^*ct*^ allele, we injected one-cell-stage *betaPix*^*ct/ct*^ embryos with *Cre* mRNA to induce global inversion of Zwitch and deletion of *betaPix*. As expected, we found *Cre*-induced precise inversion in *betaPix*^*m/m*^ as confirmed by Sanger sequencing (Figure 1C). By using *o*-dianisidine staining to label hemoglobins, we found severe brain hemorrhages developed in *betaPix*^*m/m*^ mutant with the time window similar to *bbh*^*fn40a*^ mutants (Figure 1D and 1E). Consistently, qPCR analysis showed that *betaPix* decreased and *RFP* increased in heterozygous *betaPix*^*m/+*^ and homozygous *betaPix*^*m/m*^ mutants compared with *betaPix*^*ct/ct*^ wild-type siblings, while *alphaPix* expression was not affected (Figure 1F). Of note, *TagRFP* expression in *betaPix*^*m/+*^ (after Cre-mediated recombination in *betaPix*^*ct/+*^ embryos) reflected endogenous *betaPix* expression in the boundary of midbrains and hindbrains as well as the hindbrains by light-sheet fluorescence microscopy imaging (Figure S1F and S1G). Together, these results suggest that the conditional trap *betaPix*^*ct/ct*^ zebrafish is functional for visualizing endogenous *betaPix* expression (knockin) and performing loss-of-function study (conditional knockout).

### *betaPix*^*m/m*^ mutant has brain hemorrhages, central arteries defects and abnormal glial structure that is partially rescued by Pak1 inhibitor IPA3 treatment

The main phenotypes in *bbh* mutants consist of brain hemorrhage and hydrocephalus as early as 36 hpf, as well as poor endothelial-mesenchymal contacts and defective central arteries (CtAs) sprouting in the hindbrain ^10,17^. To demonstrate the practicability of *betaPix* conditional trap lines, we assessed reported phenotypes associated with *betaPix* deficiency using homozygous *betaPix*^*ct/ct*^ zebrafish. Global *betaPix* inactivation with *Cre* mRNA injection resulted in severe cerebral hemorrhages (Figure 1D and 1E). By crossing the *betaPix*^*m/m*^ mutant line with Tg(*kdrl*:GFP) transgenic line, we found defective angiogenesis in hindbrain central arteries *via* light-sheet fluorescence microscopy imaging (Figure S2A and S2B), which is consistent with *betaPix* mutant phenotypes as reported ^10,18^.

It has been suggested that endothelial *betaPix* might not contribute to the occurrence of brain hemorrhage ^10^. During brain development, neurons and glia are important perivascular cells and they orchestrate with endothelial cells in a temporal and spatial pattern. Transgenic zebrafish that express fluorescent proteins driven by the glial fibrillary acidic protein *(gfap)* promoter have been widely used for labeling glial population ^26^. By crossing the *betaPix*^*m/m*^ mutant line with Tg(*gfap*:GFP) transgenic line, we found that radial glial structure of the hindbrain had disoriented arrangement and shorten process length in *betaPix*^*m/m*^ embryos compared to siblings by light-sheet fluorescence microscopy imaging (Figure 2A, 2B and S2C). RNA *in situ* hybridization showed that neuronal and glial precursors marker *nestin* increased while differentiated neuronal marker *pax2a* decreased at the hindbrain of *betaPix*^*m/m*^ embryos, suggesting delayed neuronal development and differentiation after *betaPix* inactivation (Figure 2C). By using global *betaPix* knockout with multiple guide-RNA simultaneously ^23^, we found that CRISPR-induced *betaPix* mutant F_0_ embryos had almost identical phenotypes to those in *betaPix*^*fn40a*^ mutants (Figure S3A and S3B), such as cerebral hemorrhages, central arteries defects, and abnormal hindbrain glial and neuronal precursor development (Figure S3C to S3G), further confirming the loss-of-function phenotypes in *betaPix*^*m/m*^mutants.

**Figure 2.**
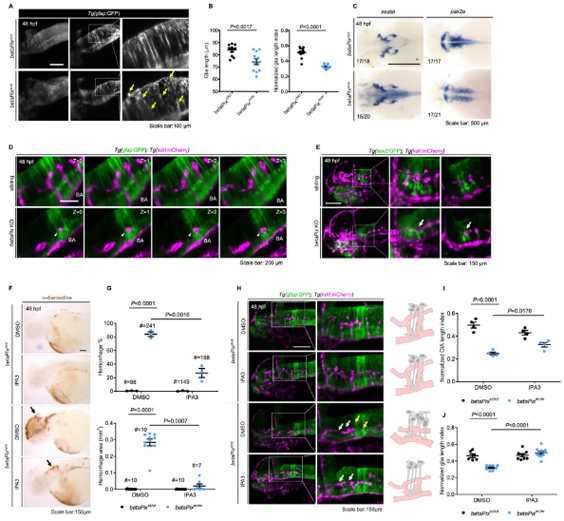
*betaPix*^*m/m*^ mutant have brain hemorrhages, central artery defects and abnormal glial structure that was partially rescued by Pak1 inhibitor IPA-3 treatment. (A) Left panel showing the maximum intensity projection of the glial structures in the hindbrain of *betaPix*^*ct/ct*^ and *betaPix*^*m/m*^ embryos at 48 hpf. Lateral view, anterior to left. Middle panel showing representative optical sections and right panel showing the higher magnifications of boxed area, presenting atypical glial structures with disoriented arrangements (yellow arrows) in *betaPix*^*m/m*^ embryos. (B) Quantification of glial parameters in (A). Left panel showing the average glia length, and right panel showing glia length index normalized to individual head length, which each dot represents one embryo. Data are presented in mean ± SEM; unpaired Student’s t test with individual *P* values mentioned in the figure. (C) Whole-mount RNA *in situ* hybridization revealed *nestin* and *pax2a* expression pattern in *betaPix*^*ct/ct*^ and *betaPix*^*mlm*^ embryos at 48 hpf. Dorsal view, anterior to the left. (D) Optical sections of glial structure (green) and blood vessels (magenta) in the heads of siblings and CRISPR-mediated *betaPix* F_0_ knockout embryos. Arteries in the hindbrain of *betaPix* KO mutants had developmental defects (white arrowheads), showing shorter distance between basilar artery and glial cell bodies. (E) 3D reconstruction of the sox2-positive precursors (green) and vasculatures (magenta) in the heads of siblings and CRISPR-mediated *betaPix* F_0_ knockout embryos at 48 hpf. Box areas are shown in higher magnifications at the middle panels, with optical sections shown in the right panels. Arrows indicate CtA with enlarged perivascular space. (F) Representative stereomicroscopy images of *o-*dianisidine staining of *betaPix*^*ct/ct*^ and *betaPix*^*m/m*^ embryos at 48 hpfthat were treated with DMSO or PAK inhibitor IPA3. Brain hemorrhages indicated with arrows. (G) Quantification of brain hemorrhagic parameters in (F). Left panel showing hemorrhage percentages, with independent experiments as dots. Right panel showing hemorrhage areas, with each dot representing one embryo, # represents the numbers of embryos scored for each analysis, and three or more individual experiments conducted. Data are presented in mean ± SEM; one-way ANOVA analysis with Dunnett’s test, individual *P* values mentioned in the figure. (H) Left panels showing 3D reconstruction of the glial structure (green) and vasculature (magenta) in the heads of *betaPix*^*ct/ct*^ and *betaPix*^*m/m*^ embryos at 48 hpftreated with DMSO or IPA3. Lateral view, anterior to left. Box areas are shown in higher magnifications at the middle panels. Defects in hindbrain central arteries indicated in white arrows, while defects in radial glia indicated in yellow arrows. Right panels showing schematic diagrams. Glia (grey) and CtAs (pink) develop normally in DMSO or IPA3-treated *betaPix*^*ct/ct*^ embryos, with fine radial glial processes and characteristic arch vasculture. Yet in *betaPix*^*m/m*^ embryos, abnormal development of the glia and central artery presented. In IPA-3-treated *betaPix*^*m/m*^ embryos, central arterial defects were partially rescued with relatively complete arch architecture, and glial processes defects significantly rescued. (I) Quantification of CtA parameters in (H). Left panel showing the average CtA length, and right panel showing the CtA length index normalized to individual head length, with each dot representing one embryo. Data are presented in mean ± SEM; one-way ANOVA analysis with Dunnett’s test, individual *P* values mentioned in the figure. (J) Quantification of glia parameters in (H). Left panel showing the average glia length, and right panel showing glia length index normalized to individual head length, which each dot represents one embryo. Individual scale bars indicated in the figure. Data are presented in mean ± SEM; one-way ANOVA analysis with Dunnett’s test, individual *P* values mentioned in the figure. Individual scale bars indicated in the figure. BA, basilar artery.

Vascularisation of the central arteries in the zebrafish hindbrain starts at 29 hpf. Tip cells start to form in the dorsal side of the primordial hindbrain channels (PHBC), sprout and migrate into the hindbrain tissue, moving towards the center around 36 hpf to establish connections with the basilar artery. A characteristic arch architecture forms at 48 hpf as boundaries dividing the hindbrain into rhombomeres^24^. In Tg(*gfap*:GFP);Tg(*kdrl*:mCherry) double transgenic embryos, CRISPR-induced *betaPix* knockouts showed that CtA sproutings were restricted to the bottom of glial processes that failed to migrate upwards to form an arched structure in compared with non-injected siblings (Figure 2D). The transcription factor SRY box-2 (Sox2) is another marker gene for neuron and glial precursors. In Tg*(sox2*:GFP); Tg(*kdrl*:mCherry) transgenic background, *Sox2*-positive precursor cells tightly wrapped around the central artery in control siblings, but *Sox2* precursor cells had loose contacts with the central arteries, with larger perivascular distances in CRISPR-induced *betaPix* mutants (Figure 2E). These results implicate the impaired interactions between endothelial cells and neuronal/glial cells during development after *betaPix* deletion.

p21-activated kinase (Pak) is a binding partner for *betaPix. Pak2a* has been shown mediating downstream signaling in *bbh*^*m292*^ mutants. Group I PAK allosteric inhibitor IPA-3 covalently modifies and stabilizes the autoinhibitory N-terminal region of PAK1, PAK2 and PAK3 ^25^. As expected, IPA-3 treatment significantly decreased incidence and intensity of cerebral hemorrhages in *betaPix*^*m/m*^ mutant embryos (Figure 2F and 2G), while IPA-3 treatment had no effect on hemorrhage induction in *betaPix*^*ct/ct*^ control siblings. In addition, central arterial defects were partially rescued in *betaPix*^*m/m*^ mutant embryos after IPA-3 treatment, showing increased endothelial protruding into the hindbrain and more arch structure formation (Figure 2H and 2I). IPA-3 treatment also decreased abnormal glial process arrangements, which the lengths of glial processes statistically reached the levels of the control siblings (Figure 2J). Thus, these data suggest that the *betaPix-Pak1/2* signaling regulates brain vascular integrity during development.

### Glial-specific *betaPix* knockouts recapture its global knockout phenotypes

To investigate *betaPix* function in different types of perivascular cells, we generated glial- or neuronal-specific transgenic lines using either *gfap* ^26^ or *huC* ^27^ promoters to drive *EGFP-Cre* fusion gene expression under *betaPix*^*ct/ct*^ background, respectively (Figure S4A). EGFP signals reported *Cre* expression while RFP signals efficiently reported Cre-induced gene trap cassette inversion and disruption of *betaPix* expressions (Figure S4B to S4F). Interestingly, glial-specific *betaPix* knockouts developed severe cerebral hemorrhages and abnormal central arteries vascularization, which were partially rescued by IPA-3 treatment (Figure 3A, 3B and 3C), highlighting *betaPix* function in glia *via* Pak1/2 signaling. On the other hand, neither vascular-, neuronal-, nor mural-specific deletion of *betaPix* had evident *betaPix* mutant phenotypes (Figure S5A to S5D). These results suggest that glial *betaPix* plays crucial roles in zebrafish embryonic vascular integrity and glial development *via* regulating *Pak* activities.

**Figure 3.**
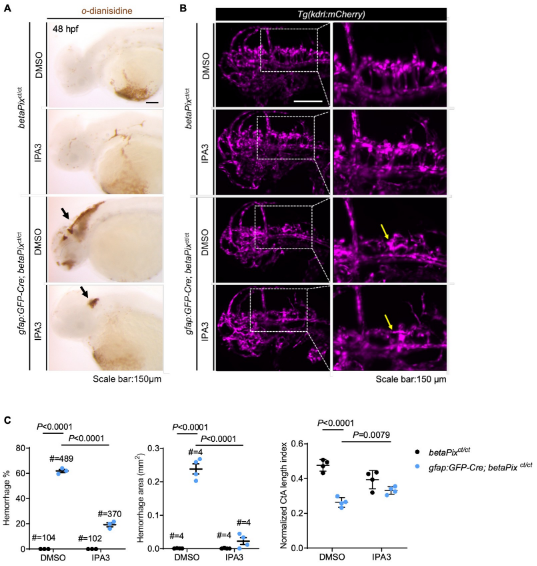
Glial-specific *betaPix* knockouts recapture global *betaPix* mutant phenotypes. (A) Representative stereomicroscopy images of erythrocytes stained with *o*-dianisidine in *betaPix*^*ct/ct*^ siblings and *gfap:GFP-Cre; betaPix*^*ct/ct*^ mutant embryos treated with DMSO or IPA3 at 48 hpf. Brain hemorrhages indicated with arrows in glial-specific *betaPix* knockouts. Lateral view with anterior to the left. (B) Left panels showing 3D reconstruction of the vasculature (magenta) in the heads at 48 hpf, lateral view with anterior to the left. Box areas are shown in higher magnifications of brain vasculatures at the right panels. CtA defects indicated in yellow arrows in *gfap:GFP-Cre; betaPix*^*ct/ct*^ mutant embryos. (C) Quantification of brain hemorrhages in (A) and CtA parameters in (B). Left panel showing hemorrhage percentages, with independent experiments as dots. Middle panel showing hemorrhage areas with each dot representing one embryo. # represents the numbers of embryos scored for each analysis, three or more individual experiments conducted. Right panel showing CtA length index normalized to individual head length, with each dot representing one embryo. Data are presented in mean ± SEM; one-way ANOVA analysis with Dunnett’s test, individual *P* values mentioned in the figure. Individual scale bars indicated in the figure.

### Single-cell transcriptome profiling reveals that *gfap*-positive progenitors were affected in *betaPix* knockouts

To investigate the interplays between glial and hemorrhagic pathology caused by *betaPix* loss-of-function, we profiled the cranial tissues of CRISPR-edited zebrafish at 1 and 2 dpf using the lOX Chromium single-cell RNA sequencing (scRNA-seq) platform (Figure 4A). Multi-guide targeting was able to generate almost 100% F_0_ null mutants, allowing us to select mutant embryos before brain hemorrhages starting from 36 hpf. Low-quality cells were excluded based on the numbers of genes detected and percentages of reads mapped to mitochondrial genes per sample. A total of 38,670 cells passed quality control and were used for subsequent analyses. Uniform manifold approximation and projection (UMAP) identified 71 cell clusters, which represented 24 zebrafish cranial cell types based on known marker gene expression profiles (Figure 4B and S6A). Enriched gene markers were compared with previously annotated gene markers in the ZFIN database and literatures^28,29^. By comparing the proportion of cells in each sample, we found that most neuronal clusters had increased relative proportions, while the numbers of glial and neuronal progenitors, endothelial cells, erythrocytes, neural crests, muscles, cartilages, retinas (photoreceptor precursor cells), olfactory bulbs, or epidermis and pharyneal arches were reduced in *betaPix* knockout heads at 2 dpf (Figure 4C and S6B). This is consistent with the open public databases of single-cell transcriptome atlas where *betaPix* is weakly expressed in a broad range of cells including glia, neurons and neuronal precursors, retinas, endothelial cells, pharynx, exocrine pancreas, olfactory cells and heart cells ^30^.Given that *betaPix* is critical in glia during brain vascular integrity development as shown above (Figures 1 and 2), we examined the *gfap* expression among each cluster, and found relatively high *gfap* expression in clusters including glial and neuronal progenitors, hindbrain, ventral diencephalon, ventral midbrain and floor plate (Figure S6A, indicated by the arrow). We next focused on the progenitor cluster owing to the enriched *gfap* expression and the significantly reduced numbers of cells in this cluster by *betaPix* deficiency. Furthermore, this progenitor cluster exhibited high-level expression of cell proliferation and cell cycle related genes *(mki67, pcna, ccndl, rrm1* and *rrm2)* as well as key glial-associated genes *(gfap, fabp7a, her4*.1, *cx43, id1, fgfbp3, atplalb* and *mdka)* (Table S1). From differentially expressed genes (DEGs) in this progenitor cluster between controls and *betaPix* knockouts at 2 dpf, gene ontology (GO) terms revealed three major categories: epigenetic remodeling, microtubule organizations, and neurotransmitter secretion/transportation (Figure 4D). Of these signaling genes, we were particularly interested in microtubule organizing genes for further studies.

**Figure 4.**
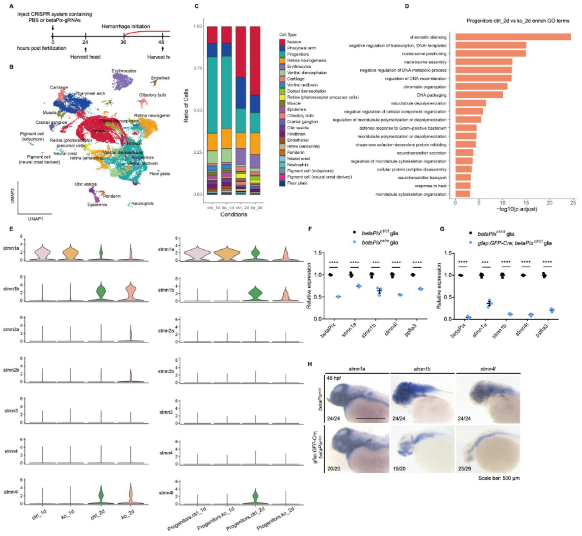
Single cell transcriptome reveals that a subcluster of glial progenitor and stathmin family members are associated with *betaPix* mutation. (A) Experimental strategy for single cell RNA sequencing of embryonic heads from wild-type siblings and *betaPix* CRISPR mutants at 1 dpf and 2 dpf. (B) UMAP visualization and clustering of cells labeled by cell type. Four samples were aggregated and analyzed together. (C) Proportions of 24 cell clusters were differentially distributed among four sample groups. ctrl_ld, PBS-injected siblings at 1 dpf; ko_ld, *betaPix* CRISPR mutants at 1 dpf; ctrl_2d, PBS-injected siblings at 2 dpf; ko_2d, *betaPix* CRISPR mutants at 2 dpf. (D) Enriched GO terms for differentially expressed genes for progenitor sub-cluster comparing ko_2d to ctrl_2d groups. (E) Violin plots showing the expression of the stathmin family genes by all cells among four sample groups (left panel) or by progenitor sub-cluster among four sample groups (right panel). (F) qRT-PCR analysis showing expression of *betaPix, stmn1a, stmn1b, stmn4l* and *ppfia3* in glia of FACS-sorted *betaPix*^*ct/ct*^ siblings and *betaPix*^*m/m*^ mutants at 48 hpf. Each dot represents cells sorted from one embryo. Data are presented in mean ± SEM. *\*\*\*P* <0.005; \*\*\*\**P*<0.001; unpaired Student’s t test. (G) qRT-PCR analysis showing expression of *betaPix, stmn1a, stmn1b, stmn4l* and *ppfia3* in glia of FACS-sorted *betaPix*^*ct/ct*^ siblings and *gfap:GFP-Cre; betaPix*^*ct/ct*^ mutants at 48 hpf. Each dot represents cells sorted from one embryo. Data are presented in mean ± SEM. *\*\*\*P* <0.005; \*\*\*\**P*<0.001; unpaired Student’s t test. (H) Whole-mount RNA *in situ* hybridization revealing down-regulation of *stmn1a, stmn1b* and *stmn4l* in *betaPix*^*ct/ct*^ siblings and *gfap:GFP-Cre; betaPix*^*ct/ct*^ mutant embryos at 48 hpf. Individual scale bars indicated in the figure.

### Stathmin acts downstream of *betaPix* in glial migration *via* regulating tubulin polymerization

Microtubules are essential cytoskeletal elements composed of α/β-tubulin heterodimers. The regulation of microtubule polymerization affects important cellular functions such as mitosis, motor transport, and migration ^31^. Based on the above scRNA-seq data (Figure 4E and S6C), we examined if microtubule-destabilizing protein Stathmins were affected in *betaPix* mutants. We found that *stathmin1a (stmn1a), stathmin1b (stmn1b*), and *stathmin4l (stmn4l)* were relatively abundant in both the whole brain and the progenitor cluster, and decreased specifically at 2 dpf in *betaPix* knockout progenitor cells (Figure 4E). Subsequent qRT-PCR revealed that these *Stathmin* expression decreased in both global *betaPix* knockouts (*betaPix*^*m/m*^) and glial-specific *betaPix* knockout glia *(gfap*:EGFP-Cre;*betaPix*^*ct/ct*^) (Figure 4F and 4G), which was further verified by RNA *in situ* hybridization analysis (Figure 4H). We next investigated whether *Stathmins* act downstream to *betaPix* signaling on regulating vascular integrity development. To this end, we found that *Pak1/2* inhibition enabled to rescue down-regulated expressions of *stmn1a, 1b* and *41* in either global *betaPix*^*m/m*^ or glial-specific *betaPix* knockout zebrafish (Figure 5A, 5B and S7A), which is consistent with previous findings that *Stathmin-1* acts downstream to *betaPix-Pak* signaling in neurite outgrowth in mice^32^.

To test whether *Stathmin* family genes are important for vascular stability, we utilized CRISPR-induced F_0_ knockout system simultaneously targeting the three stathmin genes. While singular stathmin gene knockout led to no evident phenotypes (data not shown), we found that simultaneously inactivating *stmn1a, stmn1b* and *stmn4l* caused brain hemorrhages and hydrocephalus in the brain, similar to that from *betaPix*^*fn40a*^ *and betaPix*^*m/m*^ mutants (Figure 5C, 5D, and S7B to S7E). By labeling both glia and vasculature with Tg(*gfap*:GFP; *kdrl*:mCherrry) transgenic embryos, we found that triple stathmin gene knockouts had impaired CtAs, abnormal glia and neuronal development in the hindbrain region (Figure 5E, 5F, and S7F to S7H). Furthermore, overexpressing *stmn1b* by the *gfap* promoter partially rescued brain hemorrhages and delayed neuronal development in *bbh*^*fn40a*^ mutants (Figure 5G, 5H, S7I and S7J), of which glial processes elongation also improved, but CtAs defects failed to restore (Figure 5I, 5J and S7K). Thus, these data support the critical role of *stathmins* in glia.

**Figure 5.**
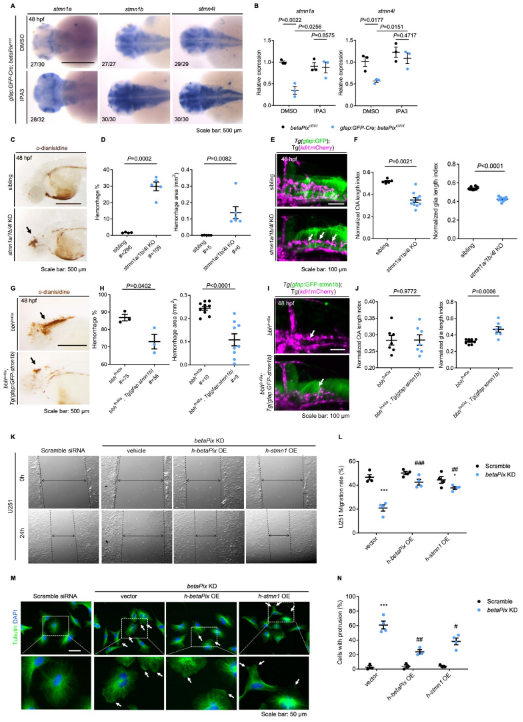
*Stathmin* acts downstream of *betaPix* in glial migration *via* regulating tubulin polymerization. (A) Whole-mount RNA *in situ* hybridization showing that *stmn1a, stmn1b and stmn4l* expression in *gfap:GFP-Cre; betaPix*^*ct/ct*^ embryos were partially rescued by Pak1 inhibitor IPA3 treatment at 48 hpf. Dorsal views, anterior to the left. (B) qRT-PCR analysis showing that *stmn1a* and *stmn4l* expression were rescued in *gfap:GFP-Cre; betaPix*^*ct/ct*^ mutants by IPA3 treatment at 48 hpf. Each dot represents one embryo. Data are presented in mean ± SEM; one-way ANOVA analysis with Dunnett’s test, individual *P* values mentioned in the figure. (C) Representative stereomicroscopy images of erythrocytes stained with *o*-dianisidine in siblings and *stmn1a/1b/4l* CRISPR mutants at 48 hpf. Brain hemorrhages, indicated with arrows, appeared in *stmn1a/1b/4l* mutants. Lateral views with anterior to the left. (D) Quantification of hemorrhagic parameters in (C). Left panel showing hemorrhage percentages, with independent experiment as dot. Right panel showing hemorrhage areas with each dot representing one embryo. # represents the numbers of embryos scored for each analysis, three or more individual experiments conducted. Data are presented in mean ± SEM; unpaired Student’s t test with individual *P* values mentioned in the figure. (E) 3D reconstruction of glial structure (green) and vasculature (magenta) in the heads at 48 hpf. Lateral view with anterior to the left. CtA defects (white arrows) indicated in *stmn1a/1b/4l* mutants. (F) Quantification of CtA and glia parameters in (E). Length index normalized to individual head length, with each dot representing one embryo. Data are presented in mean ± SEM; unpaired Student’s t test with individual *P* values mentioned in the figure. (G) Representative stereomicroscopy images of erythrocytes stained with *o*-dianisidine in *bbh*^*fn40a*^ and *bbh*^*fn40a*^; Tg*(gfap:GFP-stmn1b)* embryos at 48 hpf. Brain hemorrhages, indicated with arrows, decreased in *bbh*^*fn40a*^ mutants with glia-specific overexpression of *stmn1b*, compared with *bbh*^*fn40a*^ mutant siblings. Lateral views with anterior to the left. (H) Quantification of hemorrhagic parameters of (G). Left panel showing hemorrhage percentages, with independent experiment as dot. Right panel showing hemorrhage areas with each dot representing one embryo. # represents the numbers of embryos scored for each analysis, three or more individual experiments conducted. Data are presented in mean ± SEM; unpaired Student’s t test with individual *P* values mentioned in the figure. (I) 3D reconstruction of the *gfap:GFP-stmn1b* overexpression (green) and vasculature (magenta) in the heads of *bbh*^*fn40a*^ siblings and *bbh*^*fn40a*^; Tg*(gfap:GFP-stmn1b)* mutants at 48 hpf. Lateral view with anterior to left. White arrows indicate CtA defects. (J) Quantification of CtA and glia parameters in (I). Length index normalized to individual head length, with each dot representing one embryo. Data are presented in mean ± SEM; unpaired Student’s t test with individual *P* values mentioned in the figure. (K) Representative stereomicroscopy images of U251 cells at 0 and 24 hours after wounding. U251 cells were transfected with negative control siRNA or *betaPIX* siRNA separately, in combination with pcDNA3.1 vector, *betaPIX* overexpression plasmid, and *STMN1* overexpression plasmid. The wound edges are highlighted by dashed lines, with arrow lines indicating the wound width. (L) Quantification of wound closure in (K), showing \**P*<0.05, \*\*\**P*<0.005 compared to negative control siRNA with empty vector transfection. ^##^*P*<0.01, ^###^*P*<0.005 compared to *betaPix* knockdown with empty vector transfection. Data are presented in mean ± SEM; one-way ANOVA analysis with Dunnett’s test. (M) Representative immunofluorescence image of alpha-tubulin and DAPI signals in U251 cells. U251 cells were transfected with negative control siRNA or *betaPIX* siRNA separately, in combination with pcDNA3.1 vector, *betaPIX* overexpression plasmid, and *STMN1* overexpression plasmid. Box areas are shown in higher magnifications. Arrows indicate protrusions at the cell periphery. (N) Quantification of cell percentages with protrusions in (M). \*\*\**P*<0.005 compared to negative control siRNA with empty vector transfection. ^*#*^*P*<0.05, ^##^*P*<0.01 compared to *betaPix* knockdown with empty vector transfection. Data are presented in mean ± SEM; one-way ANOVA analysis with Dunnett’s test. Individual scale bars indicated in the figure.

To examine the direct role of *betaPix-stathmin* in glia, we turned to use human glioblastoma cell lines and manipulated the *betaPIX* levels by siRNA knockdown or transgenic overexpression. We found that *betaPIX* knockdown attenuated glial migration (Figure 5K, 5L, and S7L to S7N), which is consistent with previous studies on *betaPix* function in the endoderm ^33^, epithelial cells ^34^, neurons ^35^ and multiple carcinoma cell lines including glioblastoma ^36–38^. Either *betaPIX* or *STMNJ* overexpression led to a partial restoration of the impaired glial migration, suggesting *Stathmins* as downstream effectors of *betaPix*. Furthermore, immunofluorescence analysis revealed that *betaPIX* knockdown led to abnormal accumulation of microtubules at the cell periphery, whereas ectopic expression of *betaPIX* and *Stathmin* partially rescued these microtubule defects (Figure 5M and 5N). Together, these results suggest that *betaPix* depletion disrupts microtubule dynamics and leads to impaired glial development acting upstream to the *Pak-Stathmin* signaling, subsequently disrupting vascular integrity and resulting in brain hemorrhages. While the brain hemorrhages were only partially rescued and abnormal CtAs defects failed to restore, indicating that additional pathways may act downstream of *betaPix*.

### *Zfhx3/4* acts downstream of *betaPix* in regulating vascular integrity development

Previous studies have identified multiple signaling pathways important for zebrafish hindbrain angiogenic sprouting, such as VEGF, Notch, Wnt, Integrin β1, angiopoietin/TIE, and insulin-like growth factor signaling ^39–42^ To investigate whether angiogenic signal is disrupted by *betaPix* depletion, we examined several angiogenic gene expression pattern by performing RNA *in situ* hybridization of embryos at 36 hpf. The expression level of *Vegfaa* in sprouting CtAs decreased in *bbh*^*fn40a*^ mutants (Figure 6A). Consistently, *betaPix* knockdown significantly reduced *VEGFA* expression in cultured glial cells, which was rescued by ectopic expression of *betaPix* (Figure 6B; right panel). To explore how *betaPix* regulate *Vegf* signaling in glia, we re-examined the GO analysis of single-cell sequencing data and found a significant enrichment of epigenetic and transcriptional regulation in glial progenitor cluster (Figure 4D). Among these candidate genes, transcription factor Zinc Finger Homeobox 3 (ZFHX3) is known to mediate hypoxia-induced angiogenesis in hepatocellular carcinoma *via* transcriptional activation of VEGFA ^43^. Furthermore, ZFHX3 has been associated with stroke in multiple genome-wide association studies ^44–46^. Interestingly, we found that transcription factors *ZFHX3* and *ZFHX4* significantly downregulated after *betaPIX* inactivation in cultured glial cells (Figure 6B). The transcription levels of *Zfhx3* and *Zfhx4* also downregulated in glial progenitor cluster of our single cell RNA sequencing data upon *betaPix* knockout at 2 dpf (Figure S8A), as well as the glia-specific *betaPix* mutants (Figure S8B), thus suggesting that Zfhx3/4 might acts as downstream effectors of *betaPix*.

**Figure 6.**
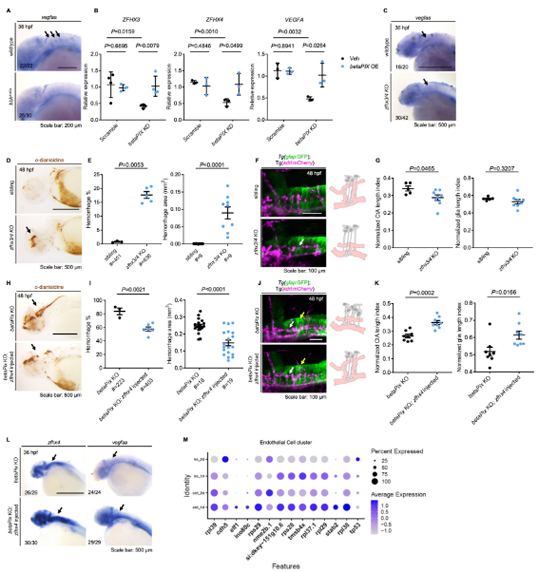
*Zfhx3/4* acts downstream of *betaPix* to regulate vascular integrity development. (A) Whole-mount RNA *in situ* hybridization revealing that *Vegfaa* decreased in *bbh*^*fn40a*^ mutants compared with siblings at 36 hpf. Lateral views with anterior to the left. Arrows indicate *vegfaa* expression in the CtAs that shown depletion in *bbh*^*fn40a*^ mutants. (B) qRT-PCR analysis revealing that *ZFHX3, ZFHX4 and* VEGFA decreased in U251 cells transfected with *betaPIX* siRNA, which were rescued by *betaPIX* overexpression. Data are presented in mean ± SEM; one-way ANOVA analysis with Dunnett’s test, individual *P* values mentioned in the figure. (C) Whole-mount RNA *in situ* hybridization showing that *vegfaa* decreased in CRISPR-mediated *zfhx3/4* F_0_ knockout embryos at 36 hpf. Lateral views with anterior to the left. Arrows indicate *vegfaa* expression in the CtAs that shown reduction in *zfhx3/4* knockouts. (D) Representative stereomicroscopy images of erythrocytes stained with *o*-dianisidine in siblings and CRISPR-mediated *zfhx3/4* F_0_ knockout embryos at 48 hpf. Arrows indicated brain hemorrhages in the knockout brains. Lateral views with anterior to the left. (E) Quantification of hemorrhagic parameters in (D). Left panel showing hemorrhage percentages, with independent experiment as dot. Right panel showing hemorrhage areas with each dot representing one embryo. # represents the numbers of embryos scored for each analysis, three or more individual experiments conducted. Data are presented in mean ± SEM; unpaired Student’s t test with individual *P* values mentioned in the figure. (F) Left panel showing 3D reconstruction of the glial structure (green) and vasculature (magenta) in the hindbrain of siblings and CRISPR-mediated *zfhx3/4* F_0_ knockouts at 48 hpf. Lateral view with anterior to the left. White arrows indicate CtA defects. Right panels showing schematic diagrams. Glia (grey) developed normally in both treatment, while defects in central artery (pink) presented in *zfhx3/4* knockout embryos. (G) Quantification of CtA and glia length parameters of (F). Length index normalized to individual head length, with each dot representing one embryo. Data are presented in mean ± SEM; unpaired Student’s t test with individual *P* values mentioned in the figure. (H) Representative stereomicroscopy images of erythrocytes stained with *o*-dianisidine in CRISPR-mediated *betaPix* FO knockout embryos with or without *zfhx4* mRNA injection at 48 hpf. Arrows indicate brain hemorrhages. Lateral views with anterior to the left. (I) Quantification of hemorrhagic parameters in (H). Left panel showing hemorrhage percentages, with independent experiment as dot. Right panel showing hemorrhage areas with each dot representing one embryo. # represents the numbers of embryos scored for each analysis, three or more individual experiments conducted. Data are presented in mean ± SEM; unpaired Student’s t test with individual *P* values mentioned in the figure. (J) Left panel showing 3D reconstruction of the glial structure (green) and vasculature (magenta) in CRISPR-mediated *betaPix* F_0_ knockout embryos with or without *zfhx4* mRNA injection at 48 hpf. White arrows indicate CtAs and yellow arrows indicate glia. Right panels showing schematic diagrams. Glia (grey) and CtA (pink) developmental defects rescued in *zfhx4* treatment. (K) Quantification of CtA and glia length parameters in (J). Length index normalized to individual head length, with each dot representing one embryo. Data are presented in mean ± SEM; unpaired Student’s t test with individual *P* values mentioned in the figure. (L) Whole-mount RNA *in situ* hybridization revealed that *Zfhx4* and *Vegfaa* decreased in CRISPR-mediated *betaPix* F_0_ knockout embryos at 36 hpf, which were rescued by *Zfhx4* mRNA injection. Lateral views with anterior to the left. Arrows indicate hindbrain regions. (M) Dot plots of several angiogenesis-associated genes expression in endothelial cell cluster. Dot size indicates the percentage of cells with gene expression, and dot color represents the average gene expression level. Individual scale bars indicated in the figure.

To determine whether *Zfhx3/4* is critical for vascular integrity by interacting with *betaPix* in zebrafish, we depleted *Zfhx3* and *Zfhx4* simultaneously *via* CRISPR-mediated Fo knockouts (Figure S8C to S8E). We found that *Zfhx3/4* knockouts decreased *Vegfaa* expression at sprouting CtAs at 36 hpf (Figure 6C). *Zfhx3/4* knockouts at 48 hpf had under-developed CtAs and reduced penetration of endothelial cells into the hindbrain (Figure 6F and 6G), developed cerebral hemorrhages during 36 to 52 hpf, which are comparable with *betaPix* mutant phenotypes (Figure 6D to 6G). Despite up-regulated expression level of *nestin*, glial arrangement and processes elongation remained statistically unaffected after *Zfhx3/4* inactivation (Figure 6F, 6G and S8F). We next investigated whether increasing expression level of *Vegfaa* in *betaPix* mutants was able to rescue impaired vascular integrity. By using *Zfhx4* mRNA microinjection into one-cell stage embryos in either *bbh*^*fn40a*^ mutants or CRISPR-induced *betaPix* knockouts, we found that ectopic expression of *Zfhx4* increased *Vegfaa* expression level (Figure 6L), thus leading to a drastic reduction of both percentages and volumes of brain hemorrhage shown in these global *betaPix* mutants (Figure 6H, 6I, and S8G to S8I), as well as improved CtAs and glial development (Figure 6J and 6K). To further explore whether angiogenic potential altered in the endothelial cells, we re-examined our scRNA-seq data and noted significant reduction of endothelial proportions among *betaPix* knockouts (Figure S6B), which showed similar trend as glial progenitors. In this endothelial cluster, CRISPR-induced *betaPix* knockouts had down-regulated angiogenic gene expression at 2 dpf (Figure 6M). Despite the lack of endothelial phenotype in cellular resolution in our present results, the disruptive transcriptome signature in endothelial cluster suggests a potential link to the defective central arterial development. Together, these data support the critical role of *Zfhx3/4* in vascular integrity and glial development probably acting upstream to VEGFA and downstream to *betaPix*.

## Discussion

Conditional knockout technology in mice has been widely used to modify and determine targeted gene function in a cell-specific manner. Spatio-temporal specific modification is beneficial for studying embryonic lethality genes or investigating the function of targeted genes in specific cell populations. Despite being broadly utilized in mice, conditional knockouts in other animal models are limited due to technical difficulties. To this end, multiple research groups have designed different strategies for establishing floxed-mutant alleles, gene traps, and inducible Cas9 in zebrafish ^47^. In particular, the Zwitch gene trap-based approach has been established successfully in zebrafish, enabling to simultaneously produce knockin reporter driven by the endogenous gene promoter and disrupting endogenous gene function in specific cell types ^19–21^. We adapted this strategy with the aid of CRISPR/Cas9 technology, and added a GSG spacer to enhance cleavage efficiency and universal guide RNA target sites on both homologous arms for producing linearized targeting vector. Moreover, we also found that utilizing either short or long homologous arms enables to achieve precise targeted integrations with this donor vector. Our short arms-based method can be broadly adapted because of less restrictions on arm sequences and easily engineering the donor vector. From the *betaPix* locus and other five unpublished loci that we already generated, we achieved Zwitch alleles with ~ 1% correct insertions based on homologous recombination as well as Cre-induced, cell-specific deletion and functional analysis of targeted genes in both embryos and adult hearts. Therefore, our CRISPR-based Zwitch method can be widely applied for generating conditional mutant zebrafish in particular and possibly extended for other animal mutants in general.

The maturation hallmark of central nervous system (CNS) vasculature is acquisition of blood brain barrier (BBB) properties, establishing a stable environment crucial for brain homeostasis in response to extrinsic factors and physiological changes. The core features of the BBB include specialized tight junctions, highly selective transporters and limited immune cell trafficking ^48,49^. In zebrafish, BBB can be functionally characterized from 3 dpf ^50,51^. *Bubblehead* mutants acquire vessel rupture phenotype early before BBB maturation, presenting a good hemorrhagic model for studying early BBB development in this work. Multiple types of cells contribute to diverse aspects of BBB development and maintenance, leading to proper vasculature function across developmental timeline. Endothelial cells form the continuous inner layer of blood vessels with junctional proteins and selective transporters, which are the main subject of permeability regulation in BBB. It is well accepted that endothelial cells do not show a predetertnined role and that the brain environment is sufficient to induce endothelial barrier ^52^. Nevertheless, accumulative evidences point to critical role of endothelial angiocrine signals in regulating several aspects of BBB development. For example, Wnt7a/b ligands secreted by neural progenitor cells bind to Frizzled receptors on endothelial cells, activating canonical Wnt signaling and downstream genes. WNT ligands knockouts lead to angiogenic defects and BBB leakages specifically in the CNS ^53^. Endothelial GPR124, as a coactivator of canonical Wnt signaling, has important function in CNS-specific angiogenesis and BBB establishment ^54–56^. Other signaling ligands such as angiopoietins and Ephrin family ligands orchestrate with Tie or EphB receptors on endothelial cells, respectively^57^.

Complex interactions between BBB cell types orchestrate proper angiogenesis and CNS development in a spatio-temporal manner. Astrocytes ensheathe capillaries through polarized end feet that is enriched with aquaporin-4 (Aqp4) proteins, colocalized with inwardly rectified K+channels^58,59^. Functional coupling of ion and water fluxes plays a critical role in regulating local osmotic equilibrium. During development, radial glia is a neuroepithelial origin with heterogeneous populations that are able to generate neurons, astrocytes, and oligodendrocytes^60–63^. In zebrafish, radial glia has long been considered serving an astrocytic role. Radial glia in zebrafish enriches with several key molecular markers for astrocytes and tight junctions while Aqp4^+^ radial glial processes rarely contact the vasculature^64^. A more recent article has reported that zebrafish G*last*-expressing radial glia transform into astrocyte-like cells, displaying dense cellular processes, tiling behavior and proximity to synapses ^65^. Whether end feet of these astrocyte-like cells enwrap capillary blood vessels and resemble mammalian astrocytes warrants future investigations. In agreement with mammalian astrocytes, two independent groups have shown that ablation of pan-glia results in progressive brain hemorrhage^66,67^ or deficiency of spinal cord arteries development in zebrafish ^68^, demonstrating important role of zebrafish glia in BBB formation. Multiple glia/astrocyte-derived signals have been shown to contribute to endothelial barrier properties development. Astrocyte-expressed sonic hedgehog (Shh) bind to hedgehog (Hh) receptors on endothelial cells and contribute to the BBB functions by promoting junctional protein expression and the quiescence of the immune system ^69–72^. Before BBB formation, notochord-derived Shh activity promote arterial cell fate on developing endothelial cells ^73^. Shh stimulates the production of angiogenic cytokines, angiopoietins and interleukins, *via* downstream transcription factor *Gli* or non-canonical pathways crosstalks with iNOS/Netrin-1/PKC, RhoA/Rock, ERK/MAPK, PI3K/Akt, Wnt/β-catenin, and Notch signaling pathways ^74^. By establishing conditional alleles of *betaPix*, we determined the critical roles of betaPix in glia, but not in other brain cells, on its cerebral vessel development and integrity. However, it remains unclear how glial betaPix is related to the above endothelial signaling to regulate endothelial cell function and blood vessel integrity.

It has been known that Pix and Pak participate in multiple signal transduction pathways ^12^. We are interested in distinctive zebrafish phenotype of the *betaPix* mutants that exhibit severe cranial hemorrhage and hydrocephalus. Previous studies have reported *betaPix-Pak2a* signaling and downstream regulation in focal adhesion essential for vascular stabilization ^10^,^17^. The experiments reported here establish glial *betaPix* mediate vascular integrity development *via* two downstream effectors (Figures 6M and 7). IPA-3 has been shown to decrease the level of phosphorylated PAK1 in a rat model of subarachnoid hemorrhage 75, and presented anti-angiogenic activity in zebrafish ^76^. Moreover, constitutively active human *Pak1* mRNA administration induced cranial hemorrhage in zebrafish ^77^. While previous researches focused on *Pak2*, our data suggest that *Pak1* also works with *betaPix* in vascular integrity development.

*Stathmin* was first discovered in a study of leukemia, of which *Stathmins are* rapidly phosphorylated when induced cells undergo terminal differentiation and stopped proliferation ^78^. Subsequent studies have showed downregulated *Stathmin* expressions after stopping proliferation of leukemia cells by chemical reagents ^79^. Similar expression changes of *Stathmins* are also found in other tumor cell lines such as breast cancer and ovarian cancer ^80,81^. *Stathmins* have also been associated with the development of the central nervous system. Systemic *Stathmin* knockout mice develop pan-neural axonopathy upon aging ^82^. Knockdown or overexpression of *Stathmin* significantly reduced the dendritic growth of Purkinje cells ^83^. Moreover, *Stathmin* is downregulated with the differentiation of oligodendrocyte progenitor cells *in vitro* ^84^. In zebrafish, *Stathmin-1* and *Stathmin-2* deficiencies resulted in defected optic nerves development with ectopic branches and axonal adhesions gaps ^85^. Knockdown or overexpression of *Stathmin-4* led to premature differentiation of dorsal midbrain neural progenitors ^86^. Systemic knockout of *Stmn4* showed cell cycle arrest in zebrafish retinal progenitor cells ^87^. These studies are consistent with our results, suggesting that transcriptionally downregulated *Stathmins* are_highly correlated with the impaired glial development in the phenotypic analysis of this study. Previous studies have revealed the role of *betaPix-d* isoform in mouse hippocampal neuron development *via* regulating microtubule stability and PAK-induced *Stathmin* phosphorylation ^32^. It remains identified whether deletion of glial- or *neuronal-betaPix* leads to brain hemorrhage. Our study found that *stathmin* knockouts impair development in a variety of cell types in the zebrafish hindbrain. Importantly, hemorrhage and hydrocephalus of *stathmin* knockouts partially phenocopy *betaPix* mutants. *In vivo* glial-specific *Stmnlb* overexpression confers partial rescue of the glial and neuronal but not vascular developmental defects in *bbh* mutants. Furthermore, ectopic expression of *betaPix* and *Stathmin* alleviate the impaired cell migration and cell shape maintenance in betaPix siRNA glioblastoma cells. Therefore, this work suggests that the *betaPix-Pak1-Stathmin* signaling regulates vascular integrity *via* glial development.

*Zfhx4* administration confers partial protection against *betaPix* depletion-induced hemorrhage, establishing the novel association between the two. However, additional components of this system remain to be discovered. It has been reported that cAMP-response element binding protein (CREB) binds to the CRE consensus site at the ZFHX3 promoter ^88^. A variety of protein kinases modulate CREB activation *via* phosphorylation at Serl 33 ^89^ including protein kinase A (PKA), phosphatidylinositol 3-kinase (PI3K)/Akt, mitogen- and stress-activated kinase 1/2 (MSK1/2), and Ca2+/calmodulin-dependent protein kinases and *Pak1* ^90–92^. Our results demonstrate that *Pak1* inhibition attenuated the majority of phenotypes induced by *betaPix* deficiency. Thus, we speculate *Zfhx3/4* are closely associated with pathogenic mechanisms downstream of *betaPix* signals by modulating expressions of angiogenetic ligands. Further investigations are required to address this hypothesis.

VEGF is the main pro-angiogenic growth factor produced by various cell types that controls multiple vascular development steps, such as proliferation, migration, permeability and survival *via* interacting with VEGF receptors on endothelial cells. Inactivation of VEGFA in astrocytes leads to BBB disruption in multiple sclerosis models ^93,94^. In early development of avascular neural tube, VEGF derived from neuroectoderm serve as a central role in inducing tip cells sprouting from the perineural vascular plexus (PVNP) and formation of vasculature lumens from stem cells ^95, 96^. Ablating neuroglia reduces *vegfab* signals and leads to ectopic intersegmental vessels sprouting and vertebral arteries in zebrafish ^68^. VEGF signaling orchestrates elaborately with other signaling pathways such as delta-like 4 (D114)/Notch signaling ^97^ and activates downstream components including the Ras/Raf/MEK, PI3K/AKT, and p38/MAPK/HSP27 pathways ^98^. Our data suggest glial *betaPix* as an important factor in vascular integrity development. Together with previous studies, *betaPix-Pak1-stathmin* signaling regulates glial growth and differentiation^32^ and *betaPix* might regulate *Vegfaa* secretion *via* downstream transcription factors *Zfhx3/4*. Thus, this work presents novel glial *betaPix* signaling in stabilizing and maintaining vascular integrity, and loss of this function causes hemorrhage and subsequent disruption of the BBB (Figure 7). However, the molecular mechanism of *betaPix* to *Zfhx3/4*, the sufficiency of *Vegfaa* and the corresponding ligand-receptor interaction between glia and endothelial cells certainly warrant future investigations.

**Figure 7.**
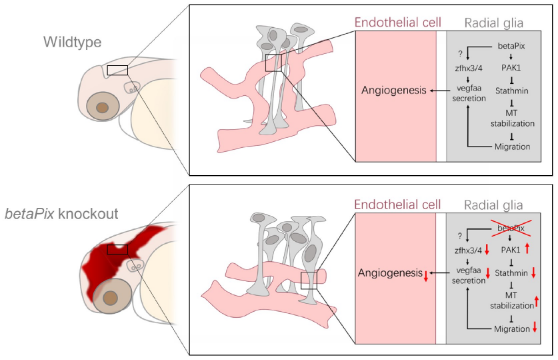
Schematic diagram on the function of glial *betaPix* in zebrafish vascular integrity development of the hindbrain. *betaPix* is enriched in glia and regulates the PAK1-Stathmin axis on microtublin stabilization, thus fine-tuning glial cell migration and their interactions with vanscular endothelial cells; and in paralelle, betaPix may also regulate Zfhx3/4-Vegfaa signaling in glia, which then modulates angiogenesis during cerebral vessel development and maturation. Deletion of *betaPix* affects glial cell migration and interaction with cerebral endothelial cells. MT, Microtubule.

Basement membrane is a non-cellular component consisting of several extracellular matrix (ECM) proteins; serve as hub for structural supports, anchoring and intercellular communication. Components of basement membrane are synthesized in surrounding cells with diverse expression patterns ^99^. For example, mice with deletion of Collagen IV in either brain microvascular endothelial cells (BMEC)- or pericytes lead to fully penetrant intracerebral hemorrhage, while COL4A1 deletion in astrocytes results in mild hemorrhage phenotypes ^100^. Differential cell-specific expression pattern of laminin isoforms contribute differently to BBB formation ^101^. As astrocytic laminins mediate pericyte differentiation and regulate capillary permeability ^102^, intercellular interactions with the ECM occur by cell surface integrin receptors. Integrin β1 on endothelial cells are critical during angiogenesis ^103^. On the other hand, integrin αvβ8 has glial-specific enrichment which mediate focal adhesion to modulate vascular integrity^11,104,105^. Notably, degradation of ECM and focal adhesions at the perivascular space is critical to the hemorrhagic phenotypes of *bbh* mutants ^17’18^. It would be of great interest to determine whether glial *betaPix* account for ECM breakdown in *bbh* mutants. Apart from *Vegfaa*, multiple pro-angiogenic growth factors and signaling ligands contribute to effective glial-vascular communication in the perivascular space. Interestingly, ZFHX3 has been reported activating a set of procollagens, integrin and platelet-derived growth factor receptor β *(Pdgfrb)* genes in response to retinoic acid stimulus, protecting cerebellar neurons from oxidative stress ^88^. In the future, it will be interesting to examine whether *Zfhx* signaling modulate adhesion molecules in the pathogenesis of *betaPix* mutants.

## Acknowledgements

The authors thank Drs. Bo Zhang, Jiulin Du, Jia Li, Liangyi Chen, Kazu Kikuchi, and Jian Chen for providing fish lines, plasmid clones, and human glial cell lines; the members of Dr. Xiong’s laboratory for helpful discussions and technical assistance; and the National Center for Protein Sciences at Peking University, particularly Dr. Liying Du at the Flow Cytometry Core for technical help on the Beckman Coulter MoFlo XDP; and Dr. Hua Liang at National Biomedical Imaging Center, Peking University for assistance with the Luxendo Multi-View Selective-Plane Illumination Microscopy (Bruker). This work is supported by grants from the National Key R&D Program of China (2019YFA0801602 and 2023YFA1800600); the National Natural Science Foundation of China (32230032 and 31730061).

## Declaration of Interest

The authors declare no competing financial interests.

## Materials and Methods

### Fish maintenance

Zebrafish were raised and handled in accordance with the guidelines of the Peking University Animal Care and Use Committee accredited by the AAALAC.

The *bbh*^*fn40a*^ mutant was isolated from a large-scale mutagenesis screen of the zebrafish genome at Massachusetts General Hospital, Boston^9^. Tg*(acta2:GFP-Cre)* was purchased from Xinjia Pharmaceutical Technology Co., Ltd (Nanjing, China); Tg*(neurod:EGFP)* were kindly provided by Dr. Bo Zhang (Peking University, China); Tg*(huC:GFP)* were kindly provided by Dr. Liangyi Chen (Peking University, China); Tg(*kdrl:GFP*) ^106^ and Tg*(kdrl:mCherry)* ^107^ were described previously.

### Cell culture and transfection

U251 and A172 human glioblastoma cell lines, kindly provided by Dr. Jian Chen at Chinese Institute for Brain Research, Beijing, were cultured in high glucose DMEM medium (Hyclone) supplemented with 10% fetal bovine serum and 1% penicillin/streptomycin, at 37°C in 5% CO_2_ humidified incubator. The *betaPIX* siRNA^37^ and *STMN1* siRNA^108^ targeting glioblastoma cell lines as previously reported were purchased from GenePharma (Shanghai, China). The sequences of the siRNAs are listed in **Table S2**. The full-length coding cDNAs of human *betaPIX* or *STMN1* were isolated from U251 cDNA library and subcloned into the pcDNA3.1 vector backbone. Transfection was conducted with Lipofectamine 3000 (Invitrogen) according to manufacturer’s instructions at a final siRNA concentration of 30 nM.

### mRNA/gRNA synthesis and microinjections

mRNA and gRNA were synthesized as described previously ^109^. In brief, linearized pT3TS-nls-zCas9-nls ^110^, *pXT7-Cre*, and *pXT7-Z*.*fhx4* plasmid DNA were purified as templates using TIANquick Mini Purification Kit (TIANGEN, DP203, China). Next, *in vitro* transcription reactions were performed using the mMESSAGE mMACHINE T3 (for pT3TS-nls-zCas9-nls) or T7 (for pXT7-Cre and *pXT7-Zfhx4)* kits (Life Technologies) according to manufacturer’s instructions. Templates for gRNA were generated by complementary annealing and elongation of two oligos. Forward oligo contained a T7 promoter and gRNA target sequence, and reverse oligo contained the universal gRNA scaffold ^23^. The resulting double-stranded DNA served as the templates for *in vitro* transcription using HiScribe® T7 High Yield RNA Synthesis Kit (NEB). The sequences of the gRNAs are listed in **Table S3**. For global *betaPix* inactivation, *Cre* mRNA with a working concentration of 200 ng/µL were injected into the *betaPix*^*ct/ct*^ embryos at one-cell stage. For CRISPR-mediated knockouts, 300 ng/µL Cas9 mRNA and 20-50 ng/µL gRNAs were injected into zebrafish embryos at one-cell stage. For *Zfhx4* rescue experiments, *Zfhx4* mRNA with a working concentration of 400 ng/µL were injected into the *bbh*^*fn40a*^ mutants or in combination with CRISPR-mediated *betaPix* knockout system at one-cell stage. Injected embryos were scored for mutant phenotypes at 36 or 48 hpf.

### Construction of *betaPix* conditional knockout zebrafish

The pZwitch plasmid clone was kindly provided by Dr. Kazu Kikuchi (National Cerebral and Cardiovascular Center Research Institute, Suita, Japan). GSG spacer was added to pZwitch+3 between P2A and TagRFP coding sequences by overlapping primer pairs. Next, a highly efficient gRNA was found from the fifth intron of *betaPix*. Primer pairs were designed to amplify the left and right homologous arms from the gRNA site for 1000 bp and 24 bp with following modifications: I) The 5’ end of the forward primer of the left homologous arm were added with an *Nhe*1 site and a universal gRNA site which provided an *in vivo* linearization cleavage site; a *Mlu*1 site to the 5’ end of the reverse primer of the left arm. 2) Added *EcoR*1 site to the 5’ end of the forward primer of the right homologous arm; added *Xho1* site and universal gRNA site to the 5’ end of the reverse primer of the right homologous arm. Then we cloned the inverted homologous arms into polyclonal sites of the modified pZwitch+3, and purified plasmid DNA with EndoFree Mini Plasmid Kit (Tiangen, DP118). *In vivo* knock-in system was composed of 300 ng/µL Cas9 mRNA, 50 ng/µL *betaPix-intron5* gRNA, 50 ng/µL Universal gRNA, and 20 ng/µL donor vector. Microinjected *in vivo* knock-in system into wildtype embryos at one-cell stage. Fish founders were screened for a-crystallin reporter expression, raised to adulthood, and re-screened for germline transmission and precise knock-in by Sanger sequencing.

### Construction of transgenic reporters

The glial reporter plasmid *gfap-EGFP* ^26^ was kindly provided by Dr Jiulin Du (Shanghai Institute for Biological Sciences, Shanghai, China). The *gfap* promoter was used to establish Tg*(gfap:GFP-Cre)* plasmid. The *stmn1b* coding sequences were amplified from wild-type zebrafish cDNA library, which were used to establish Tg*(gfap:GFP-stmn1b)* plasmid.

Transgenic reporters were generated by using To/2-based transgenesis ^111^. In brief, 100 ng/µL *Tol2* transposase mRNA in combination with 20 ng/µL donor plasmid DNA were co-injected into the embryos at one-cell stage. Transgenic founders were screened for specific transgenic GFP expression, raised to adulthood, and re-screened for germline transmission.

### Inhibitor treatment

IPA-3 (Proteintech, CM05727) were dissolved in DMSO to form a 10 mM stock, and then diluted with E3 medium to 3 µM in 6-well plates. Zebrafish embryos were treated from 24 to 48 hpf, and then washed with E3 medium for phenotypic analysis.

### *o*-dianisidine staining

*o*-dianisidine staining was performed as described previously ^18^. In brief, embryos were dechorionated, anesthetized and incubated in fresh staining solution (0.6 mg/mL *o*-dianisidine, 0.01 M sodium acetate pH4.5, 0.65% H_2_O_2_ and 40% Ethanol) for 20 minutes in dark. Washed three times with methanol, and performed benzyl alcohol/benzyl benzoate (BABB) tissue clearing before imaging with stereo microscope (Leica, 160F).

### Whole-mount *in situ* hybridization (WISH)

Whole-mount *in situ* RNA hybridization was performed as described previously ^112^. Antisense probes were synthesized using a digoxigenin RNA labeling kit (Roche, 11277073910). Primer sequences for all WISH probes used in this paper are provided in **Table S4**.

### Quantitative real-time RT-PCR

Total RNA was isolated from embryos using Trizol (invitrogen) and cDNA was generated using HiScript® III RT SuperMix (Vazyme) according to the manufacturer’s instruction. Quantitative real-time PCR was performed using Lightcycler (Roche) and ChamQ SYBR qPCR Master Mix (Vazyme). Primer sequences are listed in Supplementary Materials **Table S5**. Gene expression level was normalized against GAPDH level.

### Light-sheet fluorescence microscopy imaging

Light-sheet microscopy imaging was performed as described previously ^113^. In brief, transgenic zebrafish embryos were collected and maintained at 28.5°C. At the required developmental stages, the embryos were carefully dechorionated, paralyzed with tricaine, and transferred into 1% ultrapure low melting point agarose (16520-050; Invitrogen). The embryo was then drawn into a glass tube in top-down position using a 1 mL syringe with an 18G blunt needle. After agarose coagulation, a wire was inserted from bottom to push the agarose with embryo upwards, removed excess agar until the head area is exposed from the top of the glass tube. Finally, the glass tube was fixed in a sample holder for subsequent imaging.

Imaging was carried out with Luxendo Multi-View Selective-Plane Illumination Microscopy (Bruker). Optical calibrations were adjusted according to the instruction manual.

The optical plates are emitted from two Nikon CFI Plan Fluor 10x W 0.3 NA immersion objectives in opposite directions. The detection is completed by two Olympus 20x 1.0 NA immersion lenses. The parameters are set as follows: the green fluorescence channel uses a laser of 488 nm coupled with BP497-554 filter, the red fluorescent channel uses laser 561 nm coupled with BP580-627 filter, with an exposure time of 100 ms, a delay of 11 ms, a line mode of 50 px, and z-Stack interval of 3 µm. The raw data were stored in H5 file format, processed into TIFF format and merged Multi-View dataset using MATLAB software. After pre-processing, multi-fluorescence channels were merged in ImageJ, and 3D visualization and measurement were performed by Imaris.

### Single-cell RNA sequencing

For knockout samples, 300 ng/µL Cas9 mRNA and a mixture of number 1 to 4 *betaPix* gRNAs with 50 ng/µL each were co-injected into wildtype embryos at one-cell stage. Injection with equivalent concentration of Cas9 mRNA with PBS served as siblings. After microinjection, embryos were collected and maintained at 28.5°C. At 24 or 48 hpf, zebrafish were dechorionated, paralyzed and transferred onto agarose plate. Heads were harvested with dissecting scissors in cold sterile 1X PBS with pooling 200 heads for each group. Heads were then dissociated in 900 µL Accutase cell detachment solution (Sigma-Aldrich, A6964) at 28.5°C for 3 hours and re-suspended by pipetting every 30 min. Once digestion was complete, 100 µL FBS was added to cell suspension and centrifuged at 4°C, 500 g for 3 min. The cells were gently re-suspended in cold 500 µL 2.5% FBS in IX PBS and filtered through a 40 µm strainer. PI and Hoechst33342 were stained for distinguishing living cells.

The single, living cells were sorted by Aria SORP (BD biosciences) into 1.5 ml tubes. The cell counts and vitality were verified by AOPI staining coupled with an Automated Cell Counter (Countstar BioTech). Around 10,000-12,000 cells were loaded for each group. Single cells were barcoded with Chromium Next GEM Single Cell 3 ‘Reagent Kits V3.1 kit (10x Genomics, 1000269) in 10× Chromium Controller (10× Genomics). After qualified by peak shapes, fragment size and tailing with Fragment Analyzer System kit (Agilent Technologies, DNF-915), single-cell transcriptome libraries were sequenced via Illumina High Throughput Sequencing PE150 (Novogene, Beijing). The sequencing data were analyzed using the CellRanger-6.1.1 (10x Genomics) and mapped to reference genome GRCz1l-GRCz11.103. The output numbers of reads in four sample groups are 494,735,089 for ctrl-1d, 615,837,087 for ko-1d, 554,470,209 for ctrl-2d and 583,809,243 for ko-2d.

Low-quality cells were excluded from subsequent analyses under the following conditions: when the number of expressed genes was less than 500, when there were abnormally high counts of UMIs or genes (outliers of a normal distribution), or when the mitochondrial content exceeded 9%. A total of 38,670 cells were qualified for subsequent analyses. Unsupervised clustering was performed using Seurat (version: 4.0.2) with a resolution of 2.5, resulted in 71 cell populations and further annotated into 24 zebrafish major cranial cell types. Differential expression analysis in Seurat v4 was used to identify cluster/cell type markers by Wilcoxon rank sum test.

### Scratch assay

U251 cells were planted into culture plate and performed transfection at the desired density. At 24 hours post-transfection, we vertically scratched monolayer cells by using a 200 µL pipette tip. Washed the cells three times with PBS, then replaced with serum-free culture medium for further culture. Stereo fluorescence microscopy (Leica, 160F) was used to document the scratch size at 0 hours, 18 and 24 hours. Images were processed by ImageJ.

### ImmunostainingS

For assessing tubulin expression, U251 cells were planted into chamber slides (Saining, 1093000) and transfected at the desired density. At 24 hours post-transfection, washed the cells once on ice with pre-cooled PBS, then fixed with pre-cooled methanol at −20°C for 20 minutes. The fixed cells were permeabilized with pre-cooled acetone at −20°C for 1 minute. Removed acetone, incubated 0.5% BSA/PBS blocking solution at room temperature for 15 minutes. Cells were then incubated with 1:200 anti-a-tubulin mouse monoclonal antibody (EASYBIO, BE0031) at 37°C for 45 minutes. After rinsing once with PBS, cells were incubated with 1:400 Alexa Fluor™ 488 goat anti-mouse lgG (H+L) secondary antibody (lnvitrogen, Al 1029) at 37°C for another 45 minutes. Washed three times with PBS, and sealed with Mounting Medium with DAPI (ZSGB Bio, ZLI-9557). Images were acquired by upright fluorescence microscopy (Leica, DM5000B) at 40x magnification, and processed by ImageJ.

### Statistical analysis

Statistical analysis was performed using Graphpad Prism 6. The statistical significance of differences between the two groups was determined by the independent unpaired Student’s t-test. Among three or more groups, one-way ANOVA analysis coupled with Dunnett’s test were used. All data are presented as the mean ± SEM. *P* value <0.05 indicates significant, with individual *P* values mentioned in the figure/figure legends.

## Figure Legends

**Figure S1.**
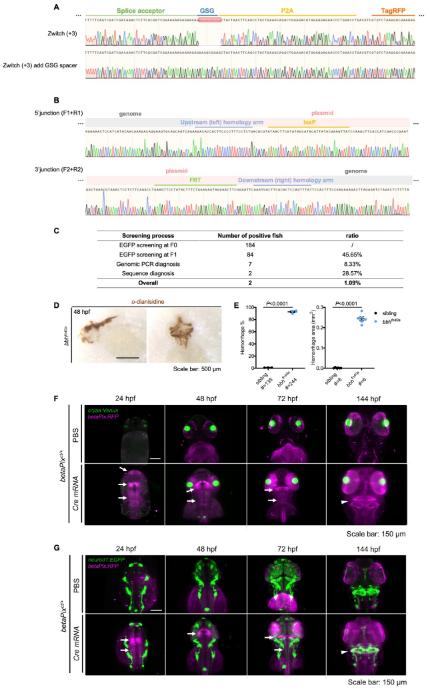
Generation of *betaPix* conditional trap (*betaPix*^*ct*^) allele by an HDR-mediated knock-in method. (A) Sanger sequencing results of the original pZwitch (+3) and the modified vector with glycine-serine-glycine spacer (GSG) in front of the 2A peptide. (B) Sanger sequencing results of the genomic PCR products at both junction sites from the *betaPix*^*ct*^ F_1_ offspring showing the correct knock-in in the betaPix locus. (C) The screening process for *betaPix*^*ct*^ knock-in zebrafish showing about 1% efficiency with correct homologous recombinations. (D) Representative stereomicroscopy images of *o*-dianisidine staining in *bbh*^*fn40a*^ embryos at 48 hpf. Lateral view, anterior to the left in left panel; Dorsal view, anterior to the top in right panel. (E) Quantification of hemorrhagic parameters in (D). Left panel showing hemorrhage percentages, with independent experiment as dot. Right panel showing hemorrhage areas with each dot representing one embryo. # represents the numbers of embryos scored for each analysis, three or more individual experiments conducted. Data are presented in mean ± SEM; unpaired Student’s t test with individual *P* values mentioned in the figure. (F) 3D reconstruction of the *betaPix* expression (magenta) in *betaPix*^*ct*/+^ brains of embryos microinjected with PBS or *Cre* mRNA at 24 hpf, 48 hpf, 72 hpf and 144 hpf. Cre-induced *RFP* expression indicates successful inversion of gene trap system (arrows). Dorsal view, anterior to the top. (G) 3D reconstruction of the *neurod1* transgenic expression (green) and *betaPix* expression (magenta) in *betaPix*^*cl/+*^ brains of embryos microinjected with PBS or *Cre* mRNA at 24 hpf, 48 hpf, 72 hpf and 144 hpf. Cre-induced *RFP* expression indicates successful inversion of gene trap system (arrows), marking strong *betaPix* expression at the midbrain hindbrain boundary (MHB), and milder expression at hindbrain at early stages. At 144 hpf, *betaPix* expresses ubiquitously in the brain, outlining the cerebellum (arrowheads). Dorsal view, anterior to the top. Individual scale bars indicated in the figure.

**Figure S2.**
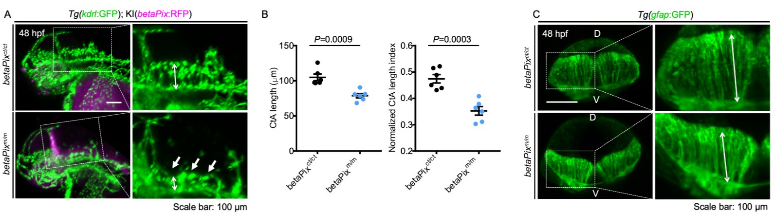
*BetaPix*^*m/m*^ mutant had brain hemorrhages, central arteries defects, and abnormal glial structure. (A) 3D reconstruction of the vasculature (green) in the heads with lateral view, anterior to left. Endogenous *betaPix* expression (magenta) only shown in Cre mRNA-injected mutant embryos. Box areas are shown in higher magnifications of vasculatures at the right panels. Central arteries (CtA) defects (white arrows) were evident in *betaPix*^*m/m*^ embryos at 48 hpf. Arrow lines indicating the CtA length. (B) Quantification of CtA parameters in (A). Left panel showing the average CtA length, and right panel showing the CtA length index normalized to individual head length, with each dot representing one embryo. Data are presented in mean ± SEM; unpaired Student’s t test with individual *P* values mentioned in the figure. (C) 3D reconstruction of the glia structure. Box areas showing higher magnifications of glia at the right panels. D, dorsal; V, ventral. Arrow lines indicating the glia length. Individual scale bars indicated in the figure.

**Figure S3.**
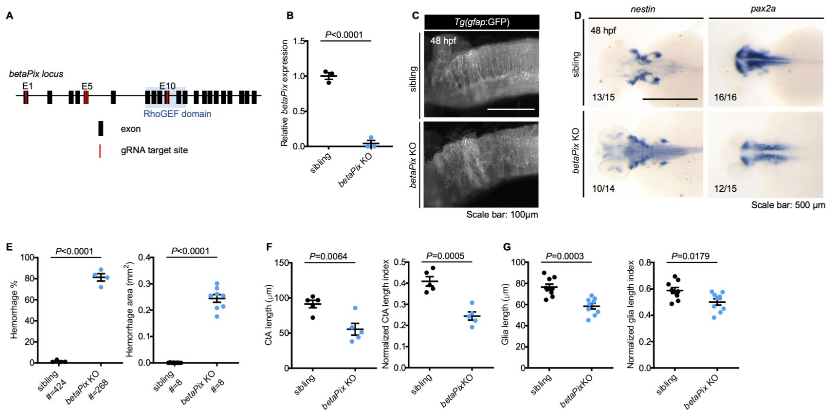
CRISPR-mediated *betaPix* F_0_ knockouts had similar phenotypes as *betaPix*^*m/m*^ mutants. (A) Schematic diagram illustrates guide RNA sites for CRISPR-mediated *betaPix* F_0_ knockout. E, exon. (B) qRT-PCR revealing that *betaPix* decreased in CRJSPR-mediated *betaPix* F_0_ knockout embryos compared with wild-type siblings at 48 hpf. Each dot represents one embryo. Data are presented in mean ± SEM; unpaired Student’s t test with individual *P* values mentioned in the figure. (C) Maximal intensity projection of the glial structure of the hindbrains in siblings and CRISPR-mediated *betaPix* F_0_ knockouts at 48 hpf, lateral view with anterior to left. (D) Whole-mount RNA *in situ* hybridization revealing up-regulated *nestin* and down-regulated *pax2a* expression in CRISPR-mediated *betaPix* F_0_ knockout embryos at 48 hpf. Dorsal view with the anterior to the left. (E) Quantification of hemorrhagic parameters of siblings and CRISPR-mediated *betaPix* F_0_ knockout embryos at 48 hpf. Left panel showing hemorrhage percentages, with independent experiment as dot. Right panel showing hemorrhage areas with each dot representing one embryo. # represents the numbers of embryos scored for each analysis, three or more individual experiments conducted. Data are presented in mean ± SEM; unpaired Student’s t test with individual *P* values mentioned in the figure. (F) Quantification of CtA parameters of siblings and CRISPR-mediated *betaPix* Fo knockout embryos at 48 hpf. Left panel showing average CtA length, andright panel showing CtA length index normalized to individual head length, with each dot representing one embryo. Data are presented in mean ± SEM; unpaired Student’s t test with individual *P* values mentioned in the figure. (G) Quantification of glial parameters of siblings and CRISPR-mediated *betaPix* Fo knockout embryos at 48 hpf. Left panel showing average glia length, and right panel showing glia length index normalized to individual head length, with each dot representing one embryo. Data are presented in mean ± SEM; unpaired Student’s t test with individual *P* values mentioned in the figure. Individual scale bars indicated in the figure.

**Figure S4.**
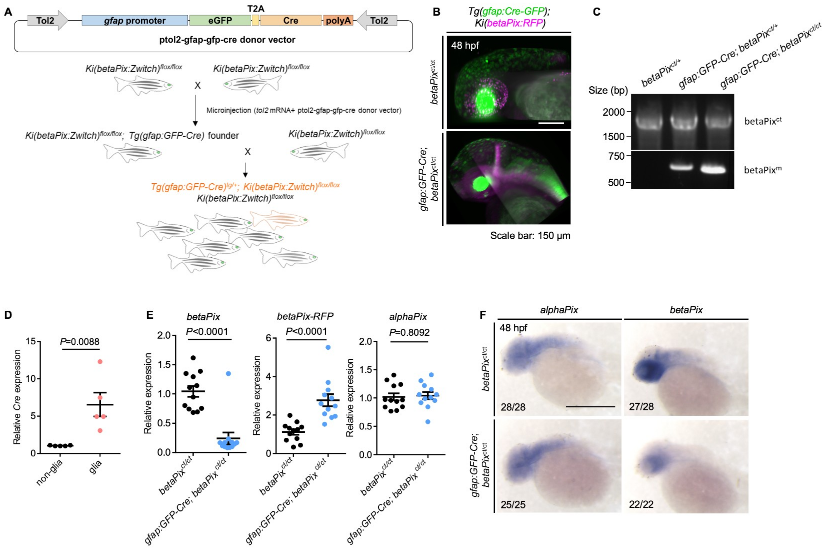
Generating glial-specific *betaPix* knockout zebrafish. (A) Schematic diagram illustrates establishment of glial-specific *betaPix* knockout zebrafish. (B) 3D reconstruction of the heads at 48 hpf with lateral view, anterior to left. Glial-specific *Cre* expression (green) was only shown in *gfap:GFP-Cre; betaPix*^*ct/ct*^ mutant embryos, which overlaps with betaPix:RFP expression (magenta). (C) Genomic PCR analysis of the *betaPix*^*ct*^ or *betaPix*^*m*^-unique sequences in *betaPix*^*ct/ct*^ siblings, *gfap:GFP-Cre; betaPix*^*ct/+*^heterozygous mutants, and *gfap:GFP-Cre; betaPix*^*ct/ct*^ homozygous mutants. (D) qRT-PCR analysis revealing *Cre* expression in *gfap:GFP-*positive glial population and *gfap:GFP-*negative non-glial population that were sorted from *gfap:GFP-Cre; betaPix*^*ct/ct*^ embryos. Each dot represents one embryo. Data are presented in mean ± SEM; unpaired Student’s t test with individual *P* values mentioned in the figure. (E) qRT-PCR analysis showing *betaPix, betaPix-RFP* and *alphaPix* expression in *betaPix*^*ct/ct*^ control siblings and *gfap:GFP-Cre; betaPix*^*ct/ct*^ mutant embryos at 48 hpf. Each dot represents one embryo. Data are presented in mean ± SEM; unpaired Student’s t test with individual *P* values mentioned in the figure. (F) Whole-mount RNA *in situ* hybridization revealing that *alphaPix* was comparable but *betaPix*^*ct/ct*^ decreased in *gfap:GFP-Cre; betaPix*^*ct/ct*^ mutants compared with *betaPix*^*ct/ct*^ siblings at 48 hpf. Lateral view with anterior to the left. Individual scale bars indicated in the figure.

**Figure S5.**
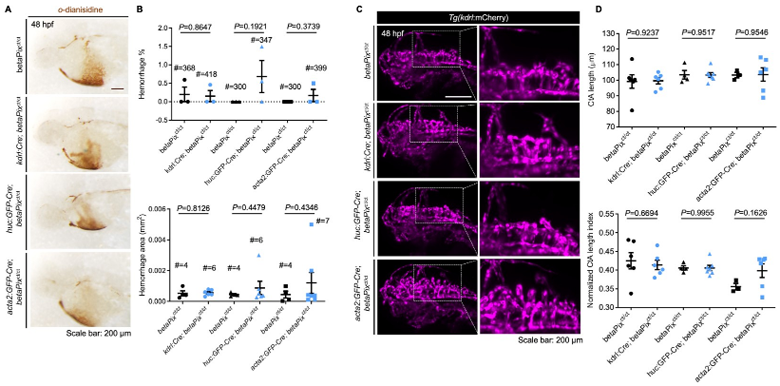
Neither endothelial-, neuronal- nor mural-specific deletion of *betaPix* caused brain hemorrhages and abnormal CtA phenotypes. (A) Representative stereomicroscopy images of *o*-dianisidine staining in *betaPix*^*ct/ct*^, *kdrl:Cre; betaPix*^ct/ct^, *huC:GFP-Cre; betaPix*^ct/ct^, and *acta2:GFP-Cre; betaPix*^ct/ct^ embryos at 48 hpf. (B) Quantification of hemorrhage parameters in (A). Up panel showing hemorrhage percentages, with independent experiment as dot. Down panel showing hemorrhage areas with each dot representing one embryo. # represents the numbers of embryos scored for each analysis, three or more individual experiments conducted. Data are presented in mean ± SEM; unpaired Student’s t test with individual *P* values mentioned in the figure. (C) 3D reconstruction of the vasculature of *betaPix*^*ct/ct*^, *kdrl:Cre; betaPix*^ct/ct^, *huC:GFP-Cre; betaPix*^ct/ct^, and *acta2:GFP-Cre; betaPix*^ct/ct^embryos at 48 hpf. Lateral view with anterior to left. Boxed areas of the hindbrains are shown in higher magnifications at the right panels. (D) Quantification of CtA parameters in (C). Up panel showing average CtA length, and down panel showing CtA length index normalized to individual head length, with each dot representing one embryo. Data are presented in mean ± SEM; unpaired Student’s t test with individual *P* values mentioned in the figure. Individual scale bars indicated in the figure.

**Figure S6.**
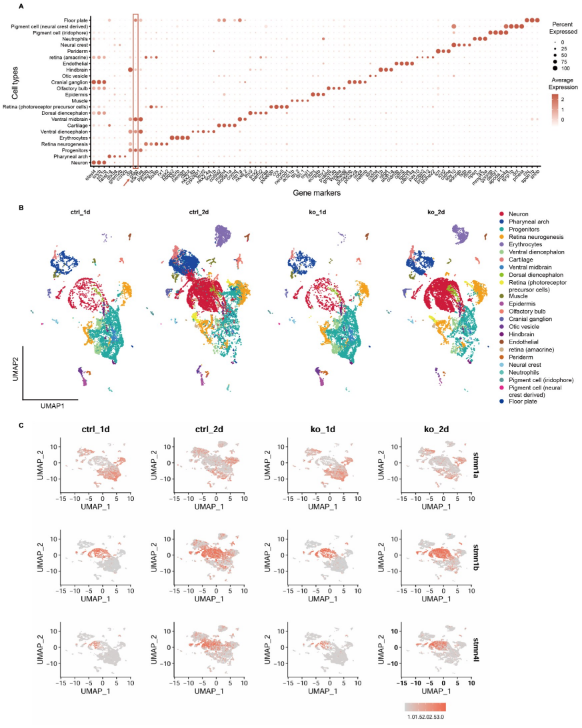
Glial progenitor sub-cluster of *betaPix* knockouts has down-regulation of *stathmin* family members and up-regulation of *pak1* gene. (A) Dot plot of marker genes expression in each cell type. Dot size indicates the percentage of cells expressed, and dot color represents the average expression level. Box area and arrow highlighting *gfap* expressions. Clusters with high *gfap* levels including neuronal and glial progenitors, hindbrain, ventral diencephalon, ventral midbrain and floor plate. (B) UMAP visualization and clustering of cells labeled by cell type across four sample groups. (C) UMAP feature plots displaying relative expression levels of selected transcripts among four sample groups. Cells are colored by expression level.

**Figure S7.**
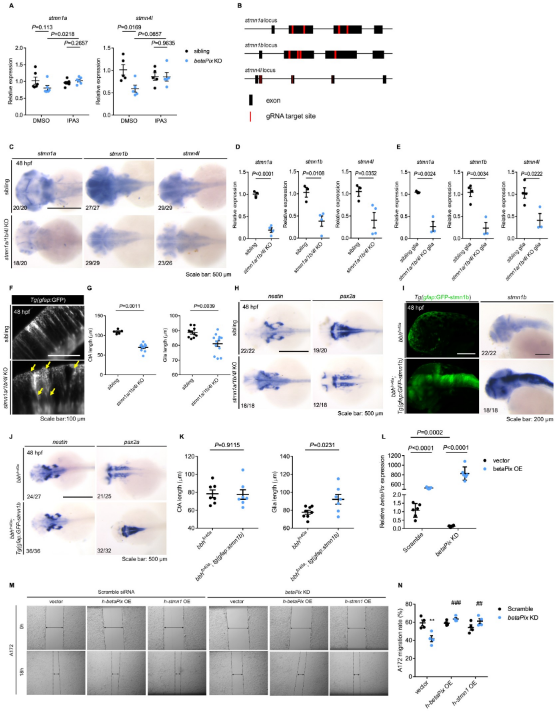
CRISPR-mediated *stmn1a/1b/4l* F_0_ knockouts have similar but milder phenotypes as *betaPix* knockouts. (A) qRT-PCR analysis showing that IPA3 treatment rescued *stmn1a* and *stmn4l* expression in CRISPR-mediated *betaPix* F_0_ knockouts at 48 hpf. Each dot represents one embryo. Data are presented in mean ± SEM; one-way ANOVA analysis with Dunnett’s test, individual P values mentioned in the figure. (B) Schematic diagram illustrates 4-guide RNAs mediated F_0_ knockout strategy in the loci of *stmn1a, stmn1b* and *stmn4l*. (C) Whole-mount RNA *in situ* hybridization confirming that *stmn1a, stmn1b* and *stmn4l* decreased in CRISPR-mediated *stmn1a/lb/41* F_0_ knockouts at 48 hpf. Dorsal views with anterior to the left. (D) qRT-PCR analysis confirtning that *stmn1a, stmn1b* and *stmn4l* decreased in mutant embryos of CRISPR-mediated *stmn1a/lb/4l* F_0_ knockouts at 48 hpf. Each dot represents one embryo. Data are presented in mean ± SEM; unpaired Student’s t test with individual *P* values mentioned in the figure. (E) qRT-PCR analysis confirming that *stmn1a, stmn1b* and *stmn4l* decreased in the FACS-sorted glia of CRISPR-mediated *stmn1a/lb/4l* F_0_ knockouts at 48 hpf. Each dot represents sorted cells from one embryo. Data are presented inmean ± SEM; unpaired Student’s t test with individual *P* values mentioned in the figure. (F) Optical sections of glial structure in the hindbrains of siblings and CRISPR-mediated *stmn1a/1b/4l* F_0_ knockouts at 48 hpf. Lateral view with anterior to the left. Abnormal glial structures with disoriented arrangements (yellow arrows) in *stmn1a/1b/4l* mutant embryos. (G) Quantification of average CtA and glial length in Figure 5 (E). Each dot represents one embryo. Data are presented in mean ± SEM; unpaired Student’s t test with individual *P* values mentioned in the figure. (H) Whole-mount RNA *in situ* hybridization revealed that *nestin* increased while *pax2a* decreased in CRISPR-mediated *stmn1a/1b/4l* F_0_ knockout embryos at 48 hpf. Dorsal views with anterior to the left. (I) 3D reconstruction and whole-mount RNA *in situ* hybridization showing that *gfap:GFP-stmn1b* transgenic overexpression rescued glial defects in *bbh*^*fn40a*^; Tg*(gfap:GFP-stmn1b)* embryos at 48 hpf. Lateral views with anterior to the left. (J) Whole-mount RNA *in situ* hybridization showing that *nestin* and *pax2a* were normally expressed in *bbh*^*fn40a*^; Tg*(gfap:GFP-stmn1b)* embryos at 48 hpf. Dorsal views with anterior to the left. (K) Quantification of average CtA and glia length in Figure 5 (I). Each dot represents one embryo. Data are presented in mean ± SEM; unpaired Student’s t test with individual *P* values mentioned in the figure. (L) qRT-PCR analysis confirming the efficacy of *betaPix* transgenic over-expression and *betaPix* siRNA knockdown in U251 cells. Data are presented in mean ± SEM; one-way ANOVA analysis with Dunnett’s test, individual P values mentioned in the figure. (M) Representative stereomicroscopy images of A172 cells at 0 and 18 hours after wounding. *betaPIX* siRNA treatment decreased A172 cell migration, which were rescued by either *betaPIX* or STMNI overexpression. The wound edges are highlighted by dashed lines, with arrow lines indicating the wound width. (N) Quantification of wound closures in (M). Data are presented in mean ± SEM; one-way ANOVA analysis with Dunnett’s test. \*\**P*<0.01 compared to negative control siRNA and empty vector transfection. ^##^*P*<0.01, ^###^*P*<0.005 compared to *betaPix* knockdown and empty vector transfecti on. Individual scale bars indicated in the figure.

**Figure S8.**
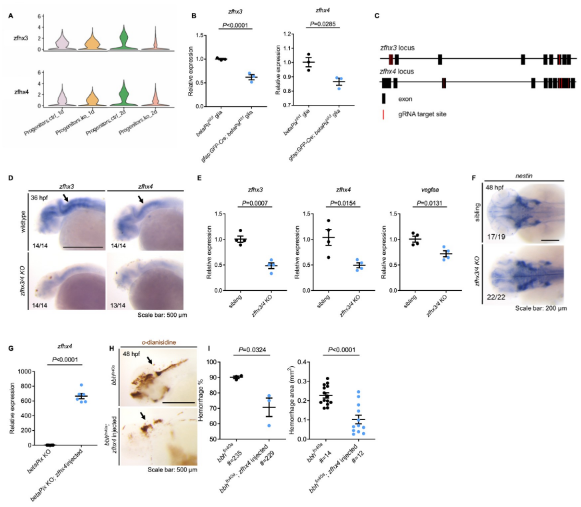
*Zfhx3/4* acts downstream of *betaPix* to regulate vascular integrity development. (A) Violin plots showing that *Zfhx3* and *Zfhx4* decreased in progenitor sub-cluster of *betaPix* knockouts. (B) qRT-PCR analysis revealing that *Zfhx3* and *Zfhx4* decreased in FACS-sorted glia of *gfap:GFP-Cre; betaPix*^*ct/ct*^ mutants compared with *betaPix*^*ct/ct*^ siblings at 48 hpf. Each dot represents the cells sorted from one embryo. Data are presented in mean ± SEM; unpaired Student’s t test with individual *P* values mentioned in the figure. (C) Schematic diagram illustrates 4-guide RNAs mediated F_0_ knockout strategy in the *Zfhx3* and *Zfhx4* loci. (D) Whole-mount RNA *in situ* hybridization confirming *Zfhx3/4* efficiency in CRISPR-mediated *Zfhx3/4* F_0_ knockout embryos at 48 hpf. Arrows indicate significant reduction of *Zfhx3* and *Zfhx4* levels in hindbrain by *Zfhx3/4* double knockout. (E) qRT-PCR analysis revealing that *Zfhx3, Zfhx4*, and *Vegfaa* decreased in CRISPR-mediated *Zfhx3/4* F_0_ knockout embryos at 36 hpf. Data are presented in mean ± SEM; unpaired Student’s t test with individual *P* values mentioned in the figure. (F) Whole-mount RNA *in situ* hybridization revealing that *nestin* increased in CRISPR-mediated *Zfhx3/4* F_0_ knockout embryos at 36 hpf. Arrows indicate the hindbrain regions. (G) qRT-PCR analysis confirming overexpression of *Zfhx4* in CRISPR-mediated *betaPix* F_0_ knockout embryos with *Zfhx4* mRNA injection at 70% epiboly. Data are presented in mean ± SEM; unpaired Student’s t test with individual *P* values mentioned in the figure. (H) Representative stereomicroscopy images of erythrocytes stained with a-dianisidine showing that brain hemorrhages (arrows) decreased in *bbh*^*fn40a*^ mutants with *Zfhx4* mRNA injection at 48 hpf. Lateral views with anterior to the left. (I) Quantification of hemorrhagic parameters in (H). Left panel showing hemorrhage percentages, with independent experiment as dot. Right panel showing hemorrhage areas with each dot representing one embryo. # represents the numbers of embryos scored for each analysis, three or more individual experiments conducted. Data are presented in mean ± SEM; unpaired Student’s t test with individual *P* values mentioned in the figure. Individual scale bars indicated in the figure.

---

Supplementary Table 1.xlsx

**Supplementary Table 2.**
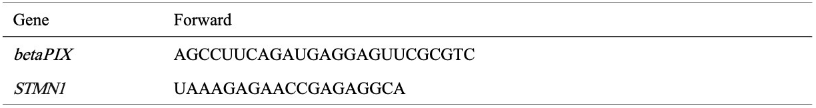
Primers used for siRNA.

**Supplementary Table 3.**
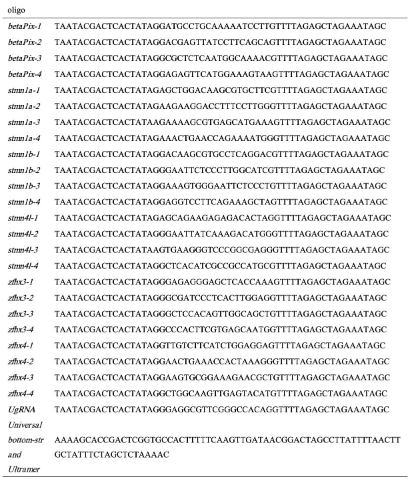
Primers used for guide RNA synthesis.

**Supplementary Table 4.**
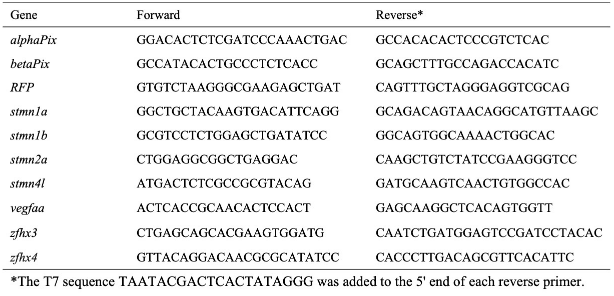
Primers used for WISH probe synthesis.

**Supplementary Table 5.**
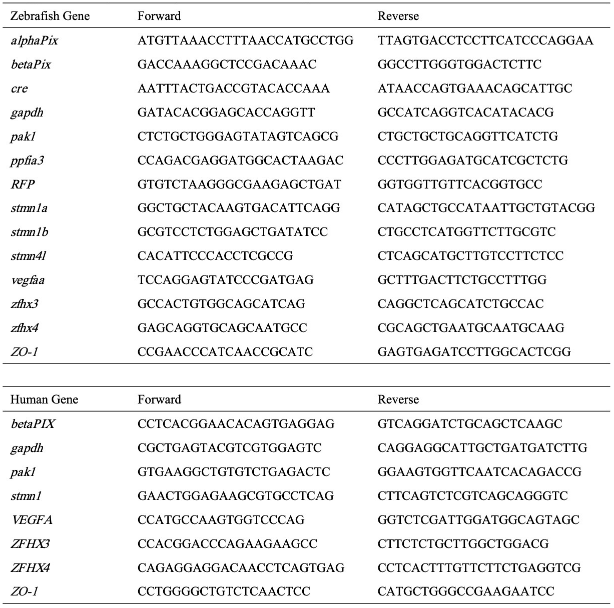
Primers used for qRT-PCR.

## Notes

https://doi.org/10.7554/eLife.106665.1

### Competing Interest Statement

The authors have declared no competing interest.

### Summary of Updates

This version is updated to correspond with the latest version submitted to eLife

